# The *PROSCOOP10* gene encodes two extracellular hydroxylated peptides and impacts flowering time in Arabidopsis

**DOI:** 10.1101/2022.09.06.506713

**Authors:** Marie-Charlotte Guillou, Thierry Balliau, Emilie Vergne, Hervé Canut, Josiane Chourré, Claudia Herrera-León, Francisco Ramos-Martín, Masoud Ahmadi-Afzadi, Nicola D’Amelio, Eric Ruelland, Michel Zivy, Jean-Pierre Renou, Elisabeth Jamet, Sébastien Aubourg

## Abstract

The Arabidopsis PROSCOOP genes belong to a family predicted to encode secreted propeptides which undergo maturation steps to produce peptides named SCOOP. Some of them are involved in defence signalling through their perception by a receptor complex including MIK2, BAK1 and BKK1. Here, we focused on the *PROSCOOP10* gene which is highly and constitutively expressed in the aerial organs. The MS/MS analyses of leaf apoplastic fluids allowed the identification of two distinct peptides, named SCOOP10#1 and SCOOP10#2, covering two different regions of PROSCOOP10. They both possess the canonical S-X-S family motif and have hydroxylated prolines. This identification in apoplastic fluids confirms for the first time the biological reality of SCOOP peptides. NMR and molecular dynamics studies showed that the SCOOP10 peptides, although largely unstructured in solution, tend to assume a hairpin-like fold exposing the two serine residues previously identified as essential for the peptide activity. Furthermore, *PROSCOOP10* mutations led to an early flowering phenotype and an increased expression of the floral integrators *SOC1* and *LEAFY*, consistent with the transcription of *PROSCOOP10* in several mutants displaying an early or late flowering phenotype. These results suggest a role of *PROSCOOP10* in flowering time, illustrating the functional complexity of the *PROSCOOP* family.

**Highlight:** The *PROSCOOP10* gene encodes two post-translationally modified extracellular SCOOP10 peptides and acts upstream of *SOC1* and *LFY* to delay flowering.

## Introduction

Small secreted peptides originate from the processing of protein precursors that share a N-terminal signal peptide addressing them to the secretory pathway. We distinguish (i) small post-translationally modified peptides (PTMPs) that are produced by proteolytic processing (Murphy *et al*., 2012); and (ii) cysteine-rich peptides (CRPs) characterised by an even number of cysteine residues involved in intramolecular disulfide bonds (Matsubayashi, 2011; Tavormina *et al*., 2015). An integrative approach combining bioinformatics, transcriptomics and phenotyping has led to the identification of a gene family encoding precursors of putative small PTMPs named Serine-riCh endOgenOus Peptides (SCOOPs) (Gully *et al*., 2019). All the predicted SCOOP peptides share a S-X-S motif and are specific to the *Brassicaceae* species. Moreover, assays based on application of synthetic peptides have shown that SCOOP peptides are phytocytokines involved in defence signalling through their perception by the MDIS1-INTERACTING leucine-rich repeat receptor kinase 2 (MIK2) (Hou *et al*., 2021; Rhodes *et al*., 2021) and the two co-receptors BRI1-associated receptor kinase (BAK1) and BAK1-LIKE 1 (BKK1) (Gully *et al*., 2019; Hou *et al*., 2021; Rhodes *et al*., 2021). New studies tolerating a greater variability in the C-terminal motif of the PROSCOOP amino acid sequence have expanded the SCOOP family to 28 members (Zhang *et al*., 2022). For example, the SECRETED TRANSMEMBRANE PEPTIDE family (STMP) contains 10 members out of which four were also annotated as SCOOP peptides: STMP1, STMP2, STMP8 and STMP10 correspond actually to SCOOP13, SCOOP14, SCOOP15 and SCOOP4 respectively (Yu *et al*., 2019). Additionally, the ENHANCER OF VASCULAR WILT RESISTANCE 1 peptide (EWR1) and four closely related peptides also encode functional SCOOP peptides (Zhang *et al*., 2022). All these peptides share the SCOOP characteristics and can induce MIK2-dependent immune responses. However, the transcription profiles of the *PROSCOOP* genes are contrasted according to organs and various stimuli. This raises the question of the involvement of the SCOOP peptides in different biological functions, notably in plant development. Indeed, *PROSCOOP12* (*AT5G44585*) is constitutively expressed in roots and recent studies have shown that SCOOP12 is a moderator of root elongation through the control of reactive oxygen species (ROS) homeostasis (Guillou *et al*., 2022). This study focuses on *PROSCOOP10* (*AT5G44580*) which is highly expressed in the aerial parts. We show that *PROSCOOP10* encodes a propeptide which gives rise to two distinct peptides that we have identified in leaf apoplastic fluids. We identify a tendency of hairpin structure of these peptides exposing the S-X-S SCOOP motif. Then, we also show an early flowering phenotype in *proscoop10* mutants suggesting a role in the development of the aerial part of the plant and particularly in flowering time.

## Materials and methods

### Plant material

*Arabidopsis thaliana* ecotype Columbia (Col-0) was used as control. Two independent *proscoop10* mutant lines in the Col-0 background were used. A first line, named *proscoop10-1*, was a T-DNA insertion line obtained from NASC (SALK_059855C) and the primers used for genotyping are listed in **Supplementary Table S1**. T-DNA insertion was checked by PCR on *PROSCOOP10* and compared with PCR on *AtCOP1*, used as a PCR positive control, in Col-0 and *proscoop10-1* mutant line. The second line, named *proscoop10-2*, was created using the CRISPR/Cas9 (clustered regularly interspaced short palindromic repeats/CRISPR-associated protein 9) method. We searched for *PROSCOOP10*-specific single guide RNA (sgRNA) and checked possible off target sites in the Arabidopsis Col-0 genome using the Crispor Tefor program (http://crispor.tefor.net). The 20-base long RNA guides with the following sequences were used: 5’-GACCACGCTCCAGGCAGTAA-3’ and 5’-ATCAGGCAGTGGGCATGGTG-3’ (**Supplementary Fig. S1A)**. Vectors and methods to get the CRISPR/Cas9 constructs were as in Charrier *et al*. (2019). Arabidopsis transformation was applied as in Zang *et al*. (2006).

The soil-grown plants used for ROS assay and phenotyping were grown under long-day conditions (16 h light at 22 °C/16 h dark at 21 °C, 70% relative humidity). Seedlings used for seedling growth inhibition assay were grown under short-day conditions (8 h light at 22 °C/8 h dark at 21 °C, 70% relative humidity). Plants used for mass spectrometry (MS) analysis were cultivated on soil under short-day conditions (8 h light at 22 °C/16 h dark at 21 °C, 70% relative humidity) during four weeks. Sodium and mercury vapor lights were used providing a light intensity of 352.9 μmol.m^-2^.s^-1^.

### Synthetic peptides

The following peptides: flg22 (QRLSTGSRINSAKDDAAGLQIA), SCOOP12 (PVRSSQSSQAGGR), SCOOP10#1 (SAIGTOSSTSDHAOGSNG), SCOOP10#2 (GDIFTGOSGSGHGGGRTOAP) were obtained from GeneCust (Boynes, France), O corresponding to hydroxyprolines. SCOOP10#2* (GDIFTGPSGSGHGGGRTPAP) corresponding to SCOOP10#2 without hydroxyprolines was obtained from Eurogentec SA (Seraing, Belgium). Peptides were synthesized with a minimum purification level of 95% and diluted in water to the final concentration used for the assays. The SCOOP10#1 and SCOOP10#2 peptide sequences were identical to the native sequences identified by mass spectrometry.

### Mass Spectrometry (MS) analyses of extracellular fluids

The extracellular fluids of rosettes were obtained according to Boudart *et al*. (2005) with slight modifications. The buffer used for the vacuum infiltration contained 5 mM sodium acetate at pH 4.6, with or without 0.3 M mannitol, and three protease inhibitors: 1 mM AEBSF (ThermoFisher, Scientific, Rockford, IL, USA), 10 mM 1-10 phenanthroline (Sigma Aldrich Chimie SARL, Saint-Quentin-Fallavier, France), and 100 μM E64 (Sigma-Aldrich). After centrifugation at 200 *g* of the vacuum-infiltrated rosettes, the fluids were collected and submitted to an ultrafiltration using an Amicon® Ultra 10K device — 10,000 NMWL (Merck Chimie SAS, Darmstadt, Germany). The samples were then speedvac-dried prior to solubilization in 10 mM DTT and 50 mM ammonium bicarbonate, followed by alkylation with 50 mM iodoacetamide. The samples were directly desalted by solid phase extraction (SPE) on C18 cartridges (StrataTM-XL 8E-S043-TG, Phenomenex, Le Pecq, France) as described in Balliau *et al*. (2018). Alternatively, the samples were subjected to tryptic digestion inside the cartridge. The peptides were eluted with 70% acetonitrile/0.06% acetic acid prior to be speedvac-dried. They were finally resuspended in 2% acetonitrile/0.1% formic acid. The samples were analysed by MS with a Q ExactiveTM-Plus Hybrid Quadrupole-Orbitrap™ mass spectrometer (Thermo-Fisher Scientific, Waltham, MA, USA) coupled with an Eksigent NanoLC-Ultra® 2D HPLC (AB SCIEXTM, Redwood City, CA, USA) as described (Balliau *et al*., 2018), except for the chromatographic separation step that was shortened to 45 min. Database search was performed as described (Duruflé *et al*., 2019), except for the enzymatic cleavage that was specified to “no enzyme”. Protein inference was performed using the X!TandemPipeline (Langella *et al*., 2017) with the following parameters: peptide E-value smaller than 0.003, protein E-value smaller than 0.01, and one peptide per protein.

### Nuclear Magnetic Resonance (NMR) analyses

SCOOP synthetic peptides were dissolved at concentrations between 0.5 mM and 1 mM in both 50 mM phosphate buffer water solution, pH 6.6, containing 10% D_2_O and in DMSO-d6. Deuterated sodium TSP-d4 at a concentration of 100 μM was used as an internal reference for chemical shift in aqueous buffers (Wishart *et al*., 1995). Measurements were performed at 278 K for aqueous and at 298 K for DMSO samples. Almost complete assignment of amide protons, non-exchanging protons and protonated ^13^C atoms was achieved in solution by ^1^H,^13^C-HSQC, ^1^H,^1^H-TOCSY (mixing time of 60 ms), and ^1^H,^1^H-NOESY (mixing time of 200 ms) recorded on a 500 MHz (11.74 T) Bruker spectrometer (Bruker France, Palaiseau) equipped with a 5 mm BBI (Broadband Inverse) probe. TopSpin 4 (Bruker BioSpin) and NMRFAM-SPARKY (Lee *et al*., 2015) were used to process and to analyse NMR data. Chemical shift deviations from random coil values were calculated using the “secondary chemical shift analysis” option of NMRFAM-SPARKY (Lee *et al*., 2015), a module based on PACSY (Lee *et al*., 2012).

### Molecular Dynamics (MD) simulations

The starting structures for our peptides were obtained by I-Tasser (Yang *et al*., 2015). *In silico* mutagenesis was performed in CHIMERA using the Rotamers tool (Pettersen *et al*., 2004). The proline residues were replaced by hydroxyproline residues using YASARA (Krieger *et al*., 2014). The GROMACS package v5.0.7 (Abraham *et al*., 2015) was used to run MD simulations. The AMBER99SB-ILDN (Lindorff-Larsen *et al*., 2010) force field was used to provide molecular mechanics parameters to our peptides. The peptides were put in a cubic cell (“box”), the border of which is at least 1 nm from the protein, and we solvate it with TIP3P explicit water molecules. Counterions were added, if necessary, to obtain a neutral system and took the place of water molecules. Energy minimization and the temperature and pressure equilibrations were done as described in Pokotylo *et al*.(2020). The equilibration time was also set at 100 ps relaxation time. Once our peptide was well-equilibrated at the desired temperature (300 K) and pressure (1 bar), we released the position restraints and run production MD for data collection. The peptides were subjected to 500 ns simulation with 2 fs time steps. To evaluate the reproducibility, the whole process (minimization, equilibration and production run) was repeated thrice. PyMol (DeLano *et al*., 2002) and VMD (Humphrey *et al*., 1996) were used for visualisation. Graphs and images were created with GNUplot (Janert *et al*., 2010) and PyMol (DeLano *et al*., 2002). All MD trajectories were analysed using GROMACS tools (Smith *et al*., 2010; Lemkul *et al*., 2018) along the last 250 ns. Polar contacts maps were determined by calculating the radial distribution function of each nitrogen and oxygen atom from all others and taking its maximum intensity in the range of H-bonds and salt-bridges. This allows to have an overview of all polar interatomic interactions and their occurrence (Ramos-Martin *et al*., 2020).

### Seedling growth inhibition assay

Seedlings were germinated on MS (1X) agar (1%) and grown for five days before transferring one seedling per well of 24-well plates containing 500 μL of MS medium or MS medium supplied with the indicated elicitor peptide to a final concentration of 1 μM (six replicates per elicitor peptide treatment). Fresh masses were measured and the experiment was repeated thrice.

### ROS assay

ROS production was determined by a luminol-based assay. Three five-week old seedlings grown on MS plates were incubated in 200 μL double distilled water (ddH_2_O) overnight in a 1.5 mL centrifuge tube. Then, ddH_2_O was replaced by 200 μL of reaction solution containing 100 μM of luminol and 10 μg/mL of horseradish peroxidase (Sigma-Aldrich, Saint-Louis, USA) supplemented with or without 1 μM peptide. Luminescence was measured for 60 min with a one-second interval, immediately after adding the solution with a FLUOstar OPTIMA plate reader (BMG LABTECH, Ortenberg, Germany). The total values of ROS production were indicated as means of the relative light units (RLUs).

### Gene expression analysis

For each of the three biological repetitions, shoot apical meristem (SAMs) samples were hand-dissected under binocular magnifier, trying to remove as much leaf tissue as possible, from 15 individual 7 and 11-day old plants of Col-0, *proscoop10-1* and *proscoop10-2*, growing under long-day conditions. Total RNAs were extracted using the Nucleospin RNA Plus Kit (Macherey-Nagel, Düren, Germany). cDNAs were synthesized from 1.5 μg of total RNA with oligo(dT) primers using Moloney Murine Leukemia Virus Reverse Transcriptase MMLV-RT according to the manufacturer’s instructions (Promega, Madison, WI, USA). RT-qPCR was carried out in a Chromo4 system (Bio-Rad, Laboratories, CA, USA). Expression profiles of key floral transition genes were calculated using the 2^−ΔΔCt^ method and were corrected as recommended in Vandesompele *et al*. (2002), with three internal reference genes (*ACT2, COP1* and *AP4M*) used for the calculation of a normalization factor. Mean expression level of Col-0 at 7 days served as calibrator. Primers for RT-qPCR analysis used are specified in **Supplementary Table S1**.

## Results

### Identification of SCOOP10 peptides in extracellular fluids by MS

Based on the high expression level of *PROSCOOP10* in leaves, we used a proteomic approach to explore the apoplastic fluid content and search for the native form(s) of SCOOP10. The experiment has been repeated thrice, twice without mannitol in the infiltration buffer and once in the presence of 0.3 M mannitol. The results were similar. In one case, a tryptic digestion of the proteins has been performed prior to the MS analysis. Different peptides could be identified covering two different regions of PROSCOOP10: they are indicated as SCOOP10#1 and SCOOP10#2 in **Fig. 1**. The corresponding MS/MS spectra are shown in **Supplementary Fig. S2**. The native SCOOP10#1 peptide comprises 18 amino acids and it covers the central predicted conserved motif defined after the comparison of the amino acid sequences of the members of the PROSCOOP family (Gully *et al*., 2019): SAIGTPSSTSDHAPGSNG (**Fig. 1A, B**). It contains the two strictly conserved serine (S) residues. All the observed peptides are native ones, as shown by the absence of tryptic sites at their N- and C-termini. SCOOP10#1 was only observed thrice compared to the 38 observations of SCOOP10#2. In all these three occurrences, it contained two hydroxyproline (O) residues. The native SCOOP10#2 peptide comprises 20 amino acids at the most and covers the C-terminal predicted conserved motif of the PROSCOOP family (Gully *et al*., 2019): GDIFTGPSGSGHGGGRTPAP (**Fig. 1A, C**). As SCOOP10#1, it also contains the two strictly conserved serine residues. It also carries three well-conserved successive glycine (G) residues. The C-terminus observed at arginine (R)16 in ten cases resulted from tryptic digestion and the corresponding peptides should not be considered as native ones. All the other observed peptides are native ones because they are not surrounded by tryptic sites. Contrary to SCOOP10#1, the pattern of proline (P) hydroxylation is variable: We could observe hydroxyproline residues at either of the three possible positions (P7, P18 or P20) and in various combinations (**Fig. 1B**). The most frequently observed position of P hydroxylation was P18 (65.8%), followed by P7 (36.8%). In some cases, P hydroxylation was observed at two positions on the same peptide: P7 and P18 (23.7%) or P7 and P20 (7.9%). The C-terminus of the peptide was variable ending with P18, alanine A19 or P20. Although protease inhibitors are used already at the beginning of the experiment, the variability of the C-terminus could be due to proteolysis by serine carboxypeptidases identified in cell wall proteomes (see *WallProtDB-2*, San Clemente and Jamet, 2015; San Clemente *et al*., 2022).

**Figure 1:**
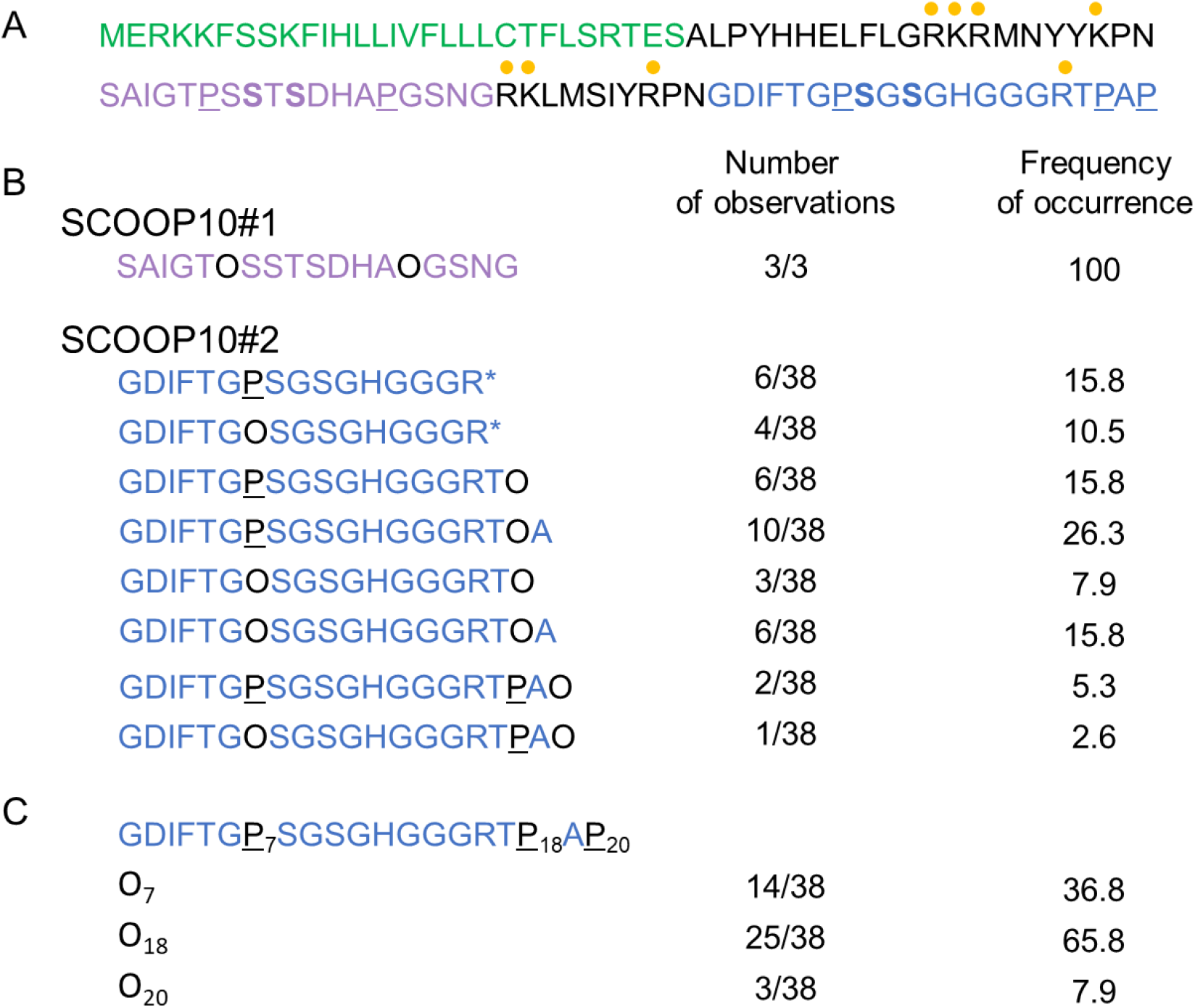
Identification of two SCOOP10 peptides by MS. **(A)** Sequence of the prepropeptide PROSCOOP10. The predicted signal peptide is in green, the mature SCOOP10#1 in purple and the mature SCOOP10#2 in blue. The tryptic cut sites (arginine (R) and lysine (K)) are indicated with yellow discs. The proline residues which were found to be hydroxylated are in black and underlined (P). The conserved serine residues are in bold. **(B)** Description of the different peptides covering the SCOOP10#1 and SCOOP10#2 amino acid sequences. The positions of P hydroxylation are indicated with O which stands for hydroxyproline. Stars indicate that the peptide has been identified after tryptic digestion. The frequency of observations of each peptide is indicated as well as the percentage of occurrences. **(C)** A focus on SCOOP10#2 to show the number of observations and the frequency of hydroxylation events at each P position.

### SCOOP10#2 is unstructured in solution but cis-trans isomerization may be favoured in a hydrophobic environment

According to MS analyses, SCOOP10#2 seems to be the major secreted or the more stable peptide produced by *PROSCOOP10*. Therefore, now knowing its exact sequence, we decided to study its structural behaviour in solution, using a hydroxylated synthetic SCOOP10#2 peptide (GDIFTGOSGSGHGGGRTOAP) and a non-hydroxylated one, SCOOP10#2* (GDIFTGPSGSGHGGGRTPAP).

NMR data indicated that the synthetic SCOOP10#2 peptide is largely unstructured in water solution. The NMR assignment of both non-hydroxylated (SCOOP10#2*) and hydroxylated (SCOOP10#2) forms are reported in **Supplementary Tables S2 and S3**, respectively. The chemical shift of some nuclei, namely Hα, Cα, Cβ and carbonyl, can be used to ascertain the presence of secondary structure by observing their deviations from random coil values (Wishart *et al*., 1992; Wishart *et al*., 1994; Wishart *et al*., 2011) which can be predicted by the peptide sequence (Nielsen *et al*., 2018). The Hα/Cα region of the ^1^H,^13^C-HSQC NMR spectrum is shown in **Fig. 2A**, while the chemical shift deviations from random coil values are reported in **Fig. 2B**. The presence of alpha helical structure can be monitored by at least three consecutive negative Hα, Cβ or positive Cα deviations while the opposite holds for beta strands. The difference between Cα and Cβ deviations is often used as a combined predictor. Overall, the observed deviations are below the threshold value of 0.1 ppm for ^1^H and 0.7 ppm for ^13^C (Wang *et al*., 2002). Hydroxylation of P7 and P18 has no major impact on the structure as indicated by minimal chemical shift perturbations on most resonances (**Fig. 2A**). Large changes are limited to the atoms of hydroxyprolines (O) and particularly to Cγ carbon atoms as expected.

**Figure 2:**
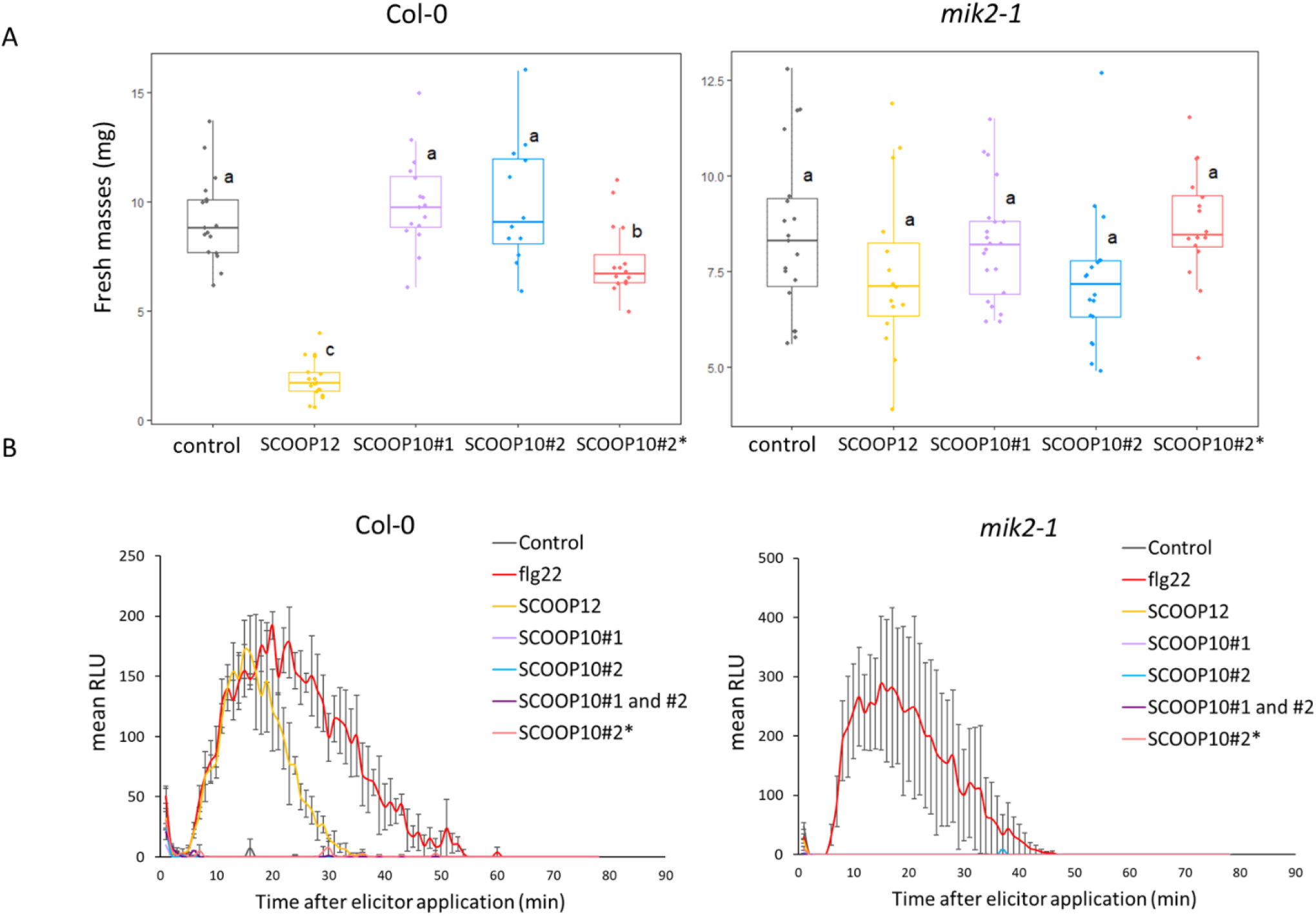
Effect of SCOOP10 peptides on seedling growth and on ROS production. **(A)** Seedling growth inhibition evaluation by fresh mass measuring after 1 μM elicitor or control treatment. **(B)** H_2_O_2_ production after 1 μM elicitor treatment or control treatment, measured with a luminol-based assay using leaf discs from 4-week-old plants of the indicated genotypes. Data represented are means of three independent replicates of over times RLU (relative luminescence units) (n =3, ±SEM). For **(A)** and **(B)**, SCOOP10#1 and SCOOP10#2 correspond to the proline hydroxylated peptides and prolines are not hydroxylated for the SCOOP10#2*peptide.

NMR spectra also allowed monitoring the conformation of the peptide bond linking proline residues to the previous amino acid (Dorman *et al*., 1973; Siemon *et al*., 1975). Such bond can often assume *cis* conformation whereas it is commonly found *trans* in all other amino acid types (Wedemeyer *et al*., 2002). A *cis* conformation determines a rather radical change in the direction of the peptide backbone and might be crucial for the biological function. In particular, the Hα of the preceding residue is close in space to the Hα or Hδ protons of proline in the *cis* and *trans* conformation, respectively. Its resonance can therefore be used to identify each conformer in NOESY or ROESY spectra (Gaggelli *et al*., 2001). A clear NOESY peak between the Hα protons of G6 and Hδ protons of P7 (or O7) revealed that the peptide bond is mainly in *trans* conformation.

In order to investigate the structural behaviour of SCOOP10#2 in a more apolar environment which could mimic that of its receptor, we studied the peptide in DMSO (**Supplementary Fig. S3 and Supplementary Tables S4 and S5**). Interestingly, we observed a doubling of peaks arising from residues 6-10, which is the region comprising the two conserved serine residues (**Fig. 1A**). Analysis of NOESY spectra reveals that the minor form might belong to the *cis* conformation of the G6-P7 peptide bond, as a cross peak is present between the Hα proton of G6 and an Hα compatible with a second form of P7. The same was not observed for P18 and P20.

### Transient hairpin-like structures exposing S8 and S10 might have functional relevance

Molecular dynamics simulations also pointed at the absence of a well-defined structure in both SCOOP10#2* and its hydroxylated form, in agreement with NMR data. Indeed, during MD simulations, an ensemble of different conformations continuously interconverted along the trajectory (**Fig. 3A**). However, these data revealed a certain tendency of the backbone to fold at the level of residues O7-S10, thus exposing S8 and S10 side chains. Interestingly, this region is the one displaying negative Hα and positive Cα-Cβ deviations in NMR, as expected for the turn of a helix (**Fig. 3B**). Although below the threshold values, these data suggest the presence of a small population of conformers contributing to the average chemical shift value. For this reason, we analysed molecular dynamic trajectories, in order to reveal key interatomic-interactions that might stabilise such a fold. For SCOOP10#2, the obtained polar contacts map reveals the formation of an H-bond between the amide nitrogen of serine S10 and the carbonyl of hydroxyproline O7 (**Fig. 3C**) or proline P7 (data not shown), a salt bridge between the side chains of aspartic acid D2 and arginine R16, and a salt bridge between the C- and N-termini (**Fig. 3D)**. Furthermore, a stacking between the aromatic rings of phenylalanine F4 and histidine H12 might also further stabilise the conformation (**Fig. 3D**). Similar contacts are observed in the non-hydroxylated form (data not shown).

**Figure 3:**
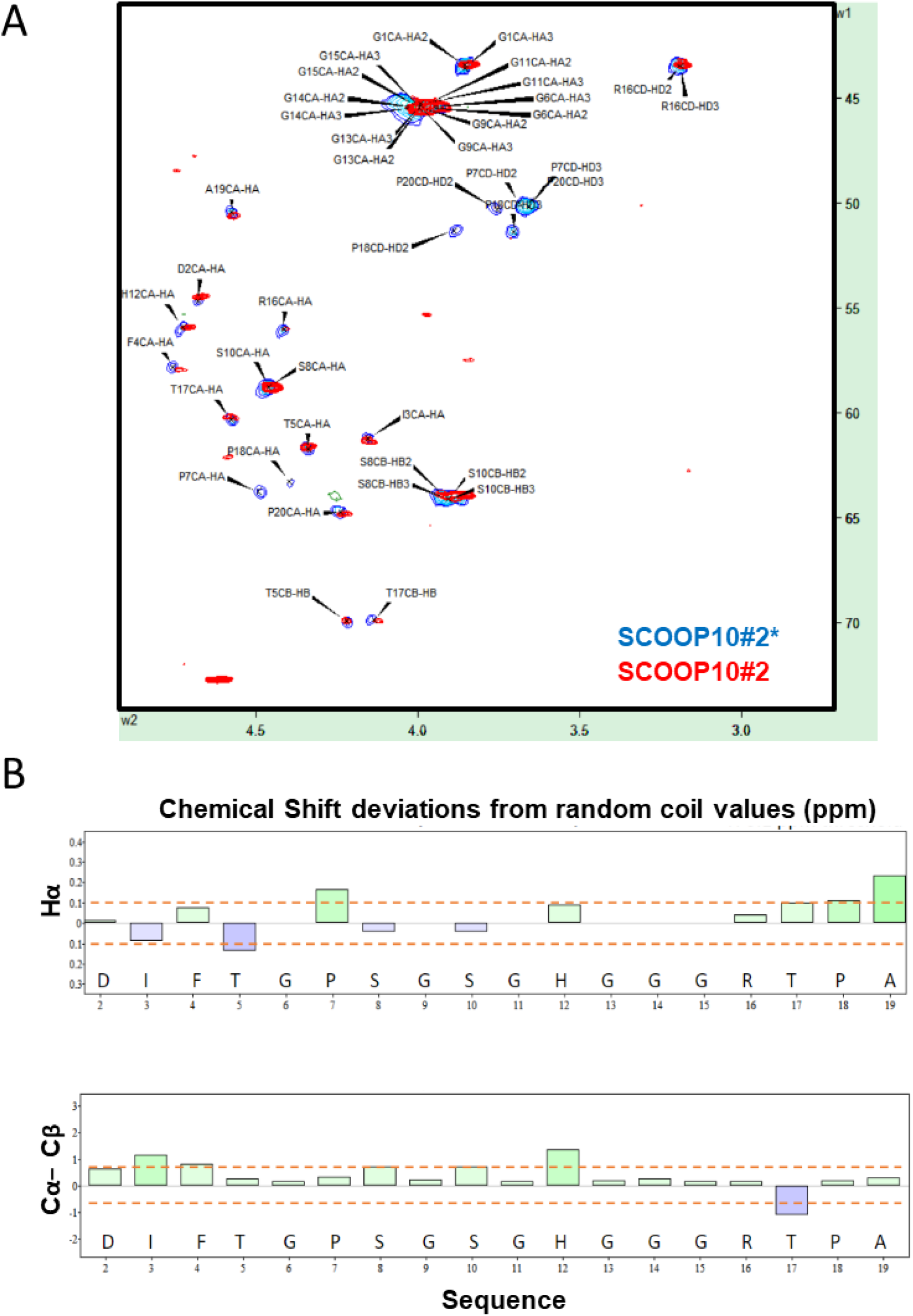
Structural behaviour of SCOOP10#2 and SCOOP10#2* in solution. **(A)** ^1^H,^13^C-HSQC spectrum assignment of non-hydroxylated SCOOP10#2* (blue) and hydroxylated SCOOP10#2 (red) 0.5 mM in 50 mM phosphate buffer at pH 6.6 and 278 K. **(B)** Chemical shifts deviations from random coil values of Hα protons and of the difference between Cα and Cβ carbons suggest the absence of a well definite structure for SCOOP10#2. Deviations for glycine Hα atoms were intentionally omitted.

In order to test the relative contribution of each of these key inter-residue interactions we performed multiple molecular dynamic simulations where we mutated at least one partner residues: R16A, G13,14,15A, P7A and S8,10A (**Supplementary Figure S4A**). In the R16A mutant, where we eliminated the salt bridge between D2 and R16, SCOOP10#2* still tends to fold onto itself but at the level of residues 12-15 (**Supplementary Figure S4B**), thus reducing the exposition of S8 (**Supplementary Figure S5**). In this case the driving force is the salt bridge between the N and C termini. When glycine residues in the flexible region G13,14,15 are mutated to alanine, the N-C terminal interaction favours a fold in the middle of the structure (residues 9-10) (**Supplementary Fig. S4B**). As for the H-bond between the amide of S10 and P7, mutants (P7A and double mutant S8,10A) reveal small perturbation of the structural behaviour (in P7A the structure folds around residue 8 by a turn rather than a helix 3-10), an effect somehow expected considering that the interaction is established at the level of the backbone.

### SCOOP10#1 is unstructured in solution with transient head-to-tail contact

We also studied the structural behaviour in solution of SCOOP10#1, using a synthetic SCOOP10#1 peptide identical to the native form identified (SAIGTOSSTSDHAOGSNG). For the synthetic SCOOP10#1 peptide, deviations from random coil values also indicate a poor structuring (**Supplementary Fig. S6**, and the NMR assignment in different conditions in **Supplementary Tables S6 and S7**), and NOESY spectra are compatible with *trans* conformation for the peptide bonds involving both P6 and P14. Contrarily to what observed for SCOOP10#2, we could not detect the presence of *cis* conformation in the more apolar environment of DMSO. MD simulations (**Supplementary Fig. S7**) agree with NMR data and detect lack of structure (**Supplementary Fig. S7A**), however, infrequent contacts between the N- and C-termini (**Supplementary Fig. S7B**) generate folded conformations vaguely resembling those observed for SCOOP10#2 (see contact map in **Supplementary. Fig. S7C)**.

### SCOOP10 peptides do not induce ROS production and growth inhibition

Effects of the SCOOP10 synthetic peptides on seedling growth and ROS production were tested on Col-0 and *mik2-1* genotypes and compared to those of the SCOOP12 peptide, as a control (**Fig. 4**). The SCOOP12 amino acid sequence was the same already used by Gully *et al*. (2019) and Guillou *et al*. (2022) and based on prediction without post-translational modifications.

**Figure 4:**
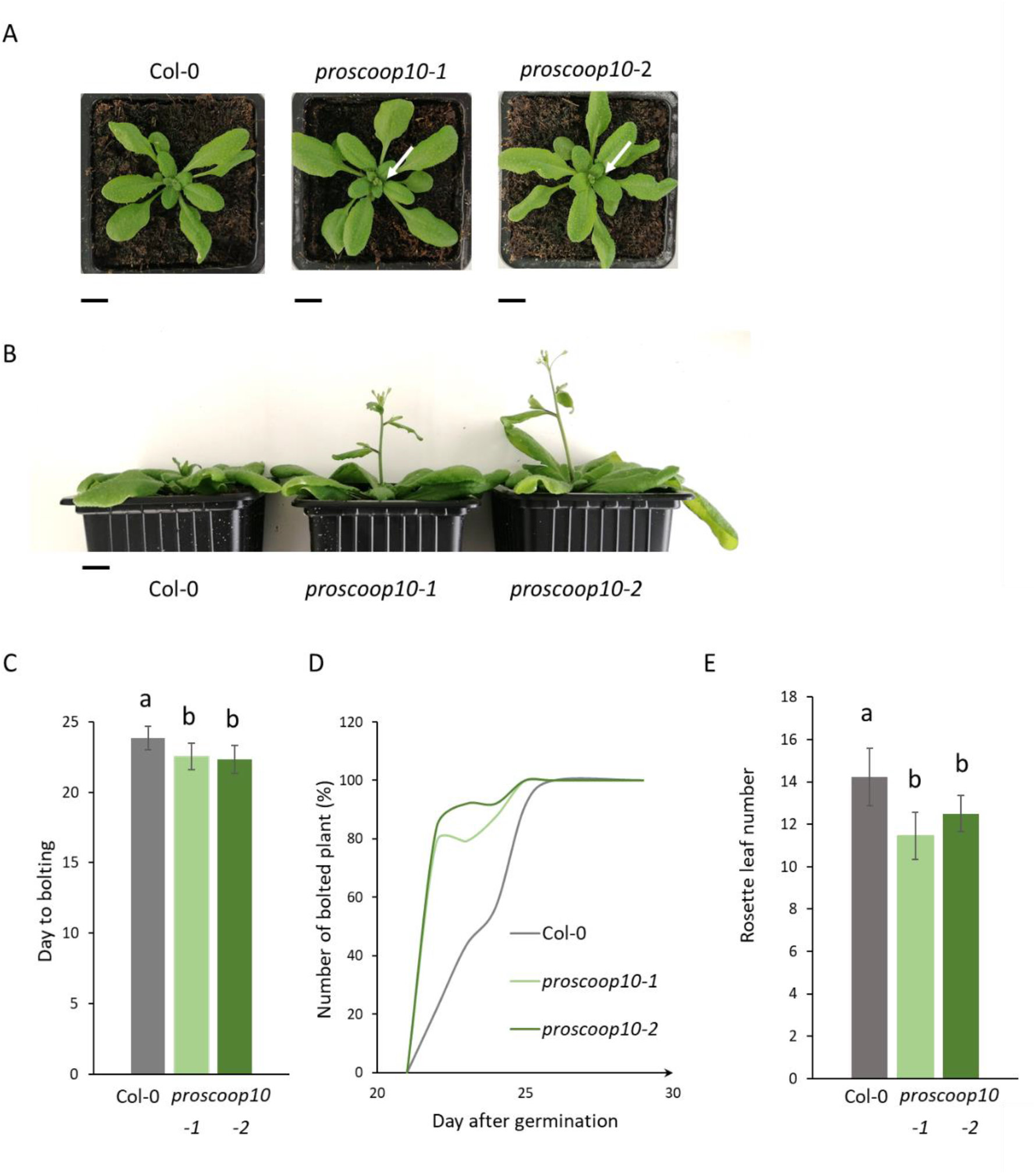
*proscoop10* early flowering phenotype. (**A**) Bolted inflorescence of the *proscoop10* mutant lines, the 22^nd^ day after germination, indicated by the white arrows. (**B**) Delay of stem development between mutants and WT due to the early flowering of the *proscoop10* lines, at the 27^th^ day after germination. Scale bar = 1cm for (**A**) and (**B**). (**C**) Average of day to bolting for each genotype. (**D**) Kinetics represent the number of bolted inflorescences in %, starting from day 20 to day 30 after germination. (n=25). (**E**) Average of the number of rosette leaves at the flowering time for each genotype. For (**C**) and (**E)** ANOVA and Tukey test allowed to define significantly two different groups labelled a and b.

In Col-0 background and at 1 μM, we showed that, the SCOOP10#2* peptide induced a seedling growth inhibition which was much smaller than that induced by the SCOOP12 peptide. In the same conditions, the SCOOP10#1 and SCOOP10#2 peptides did not show a significant effect. In the *mik2-1* mutant background, none of the SCOOP peptides had an effect on growth, as expected (**Fig. 4A**). At the same concentration, the SCOOP12 peptide induced a strong ROS production in Col-0 leaves, as already reported (Gully *et al*., 2019; Rhodes *et al*., 2021). On the contrary, neither the SCOOP10#2* nor the SCOOP10#1 and the SCOOP10#2 peptides, even simultaneously applied, induce ROS production in leaves. As expected none of the SCOOP peptides induced a high ROS production in the *mik2-1* mutant (**Fig. 4B**).

### Mutations of *PROSCOOP10* impact flowering time

Previous analysis of transcriptomic data showed that *PROSCOOP10* is highly expressed in shoot apex and leaves and may play a role in aerial organ development (Gully *et al*., 2019; Hou *et al*., 2021). Therefore, we compared the leaf development and flowering time of wild-type and *proscoop10* plants. The first *proscoop10* mutant (*proscoop10-1*) is a T-DNA insertion line ordered from NASC (**Supplementary Fig. S1A**). This line was genotyped as mutated on both DNA and cDNA (**Supplementary Fig. S1B**). We generated a second line (*proscoop10-2*) using a CRISPR/Cas9 approach and the mutant was genotyped (**Supplementary Fig. S1C, D**). *proscoop10-2* was a bi allelic mutant, which is frequently observed with this approach (Pauwels *et al*., 2018). In the first modified allele, a deletion of 237 bp occurred between the two guides in addition to modifications of nucleotides within the guides, leading to a frameshift. For the second modified allele, a deletion of 448 bp occurred between the two guides (**Supplementary Fig. S1E**). In each case, the modifications prevented the synthesis of the native SCOOP10 peptides (**Supplementary Fig. S1F**).

Both mutant lines displayed a normal vegetative development but 22 days after germination, the inflorescences of *proscoop10* mutants bolted significantly earlier than those of Col-0 plants, with a shift of two days (**Fig. 5A, C**) and the number of rosette leaves were significantly lower for *proscoop10* at bolting day (**Fig. 5E**). Consequently, five days after, mutants had longer stem length compared to Col-0 (**Fig. 5B**). 80% of the inflorescence of *proscoop10* mutants bolted around the 22^nd^ day after germination whereas the inflorescence of wild-type bolted gradually from the 23^rd^ to the 26^th^ day after germination (**Fig. 5D, Supplementary Table S8**).

**Figure 5:**
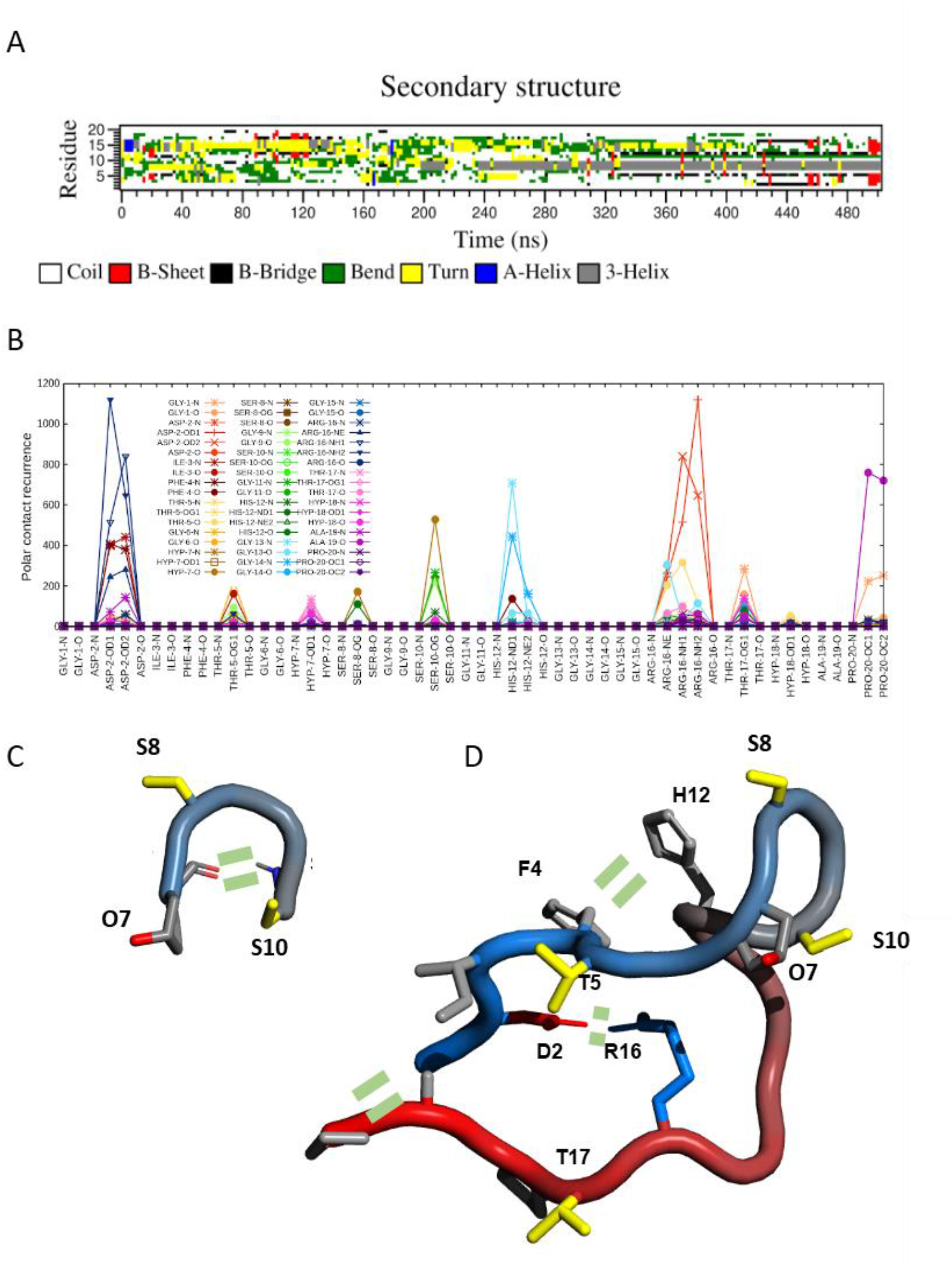
Secondary structures and intramolecular interactions found in MD simulations of SCOOP10#2. **(A)** DSSP secondary structures calculated along molecular dynamics (MD) simulation of hydroxylated SCOOP10#2 in solution; **(B)** Occurrence of intramolecular polar atom contacts (H-bonds and salt bridges) in SCOOP10#2 calculated along MD simulation trajectories; **(C, D)** Schematic representations of SCOOP10#2 shown as a ‘tube’ coloured from blue (N-terminus) to red (C-terminus). Side-chains are shown as sticks with the following color code: positively charged (blue) and non-polar (light gray). The structures were created with PyMol (DeLano et al., 2002). Key intramolecular interactions are indicated by two short parallel dashes.

Based on the mutant phenotype, we next examined the expression of three genes involved in floral regulation, *SUPPRESSOR OF CONSTANS 1* (*SOC1), LEAFY* and *GA3OX-1*, in the SAM of *proscoop10* mutants and Col-0 at 7 and 11 days after germination (**Figure 6**). At 7 days, the two floral transition genes, *SOC1* and *LEAFY*, were not differentially expressed between mutants and Col-0 whereas *GA3OX-1* was induced only in *proscoop10-1*. At 11 days, all three genes were up-regulated in the two *proscoop10* mutant lines compared to Col-0.

**Figure 6:**
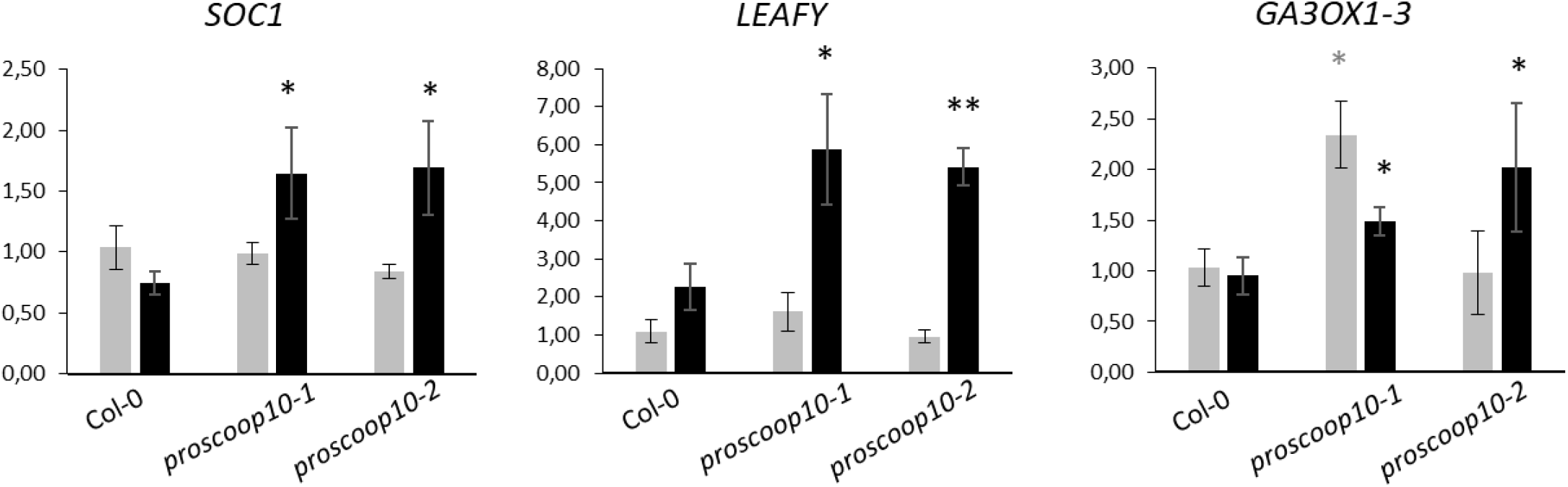
Impact of *PROSCOOP10* mutation on transcription of genes involved in floral regulation. Relative expression of *SOC1, LEAFY* and *GA3OX-1* genes was measured by RT-qPCR after 7 days (in grey) and 11 days (in black) in the two *proscoop10* mutant lines compared to Col-0 at 7 days. Values represent mean ratios ± SEM of three independent biological replicates. Asterisks denote statistical differences of gene expression levels between *prosccoop10* mutants and Col-0: \**P* < 0.05, \*\**P* < 0.01 (ratio paired t-test).

## Discussion

### Two distinct hydroxylated SCOOP10 peptides are present in the leaf apoplasm

The MS analysis of apoplastic fluid samples from rosette leaves has identified two hydroxylated SCOOP10 peptides (18 and 20 aa-long) resulting from the processing of two distinct regions of the PROSCOOP10 protein separated by 10 amino acids. This result confirms that *PROSCOOP* genes are indeed encoding preproproteins processed in short secreted peptides. Furthermore, the presence of hydroxyproline residues confirms that SCOOPs are PTMPs according to the Matsubayashi’s classification (Matsubayashi, 2011). We thus characterized two distinct SCOOP10 peptides showing the biological reality of the previous predicted peptides SCOOP10#1 and SCOOP10#2. Note that the observed native peptides are a few amino acids longer (at both their N- and C-termini) than previously predicted. As shown by MS analyses, SCOOP10#2, corresponding to C-terminal end of the PROSCOOP10 precursor, seems to be the major form in the leave apoplasm. The ability of a precursor protein to be processed in different peptides has been described for a few genes belonging to CLE, CEP and PIP families (Murphy *et al*., 2012; Roberts *et al*., 2013; Vie *et al*., 2015). In our case, the ability of *PROSCOOP10* to produce two distinct SCOOP10 peptides probably comes from the local duplication of an exon. Indeed, contrary to the large majority of *PROSCOOP* genes which have two coding exons (the first one encoding the signal peptide, and the second one containing the conserved SCOOP motif), *PROSCOOP10* has a third exon containing a second SCOOP motif (**Supplementary Fig. S1A**, Gully *et al*., 2019). This feature is also shared by *PROSCOOP6, 7, 11* and *15* which are also probably able to encode two SCOOP peptides even if previous assays based on exogenous application of predicted synthetic peptides showed that only the C-terminal ones have a biological activity (Hou *et al*., 2021; Rhodes *et al*., 2021). The identification of these native SCOOP10 peptides suggests that their maturation requires cleavage steps by endoproteases still unknown. The N-ter termini of both SCOOP10 peptides are located upstream the **Y**[KR]**P**N motif (**Figure 1**) similar to the cleavage site of IDA precursors where P and Y residues in positions −2 and −4 relative to the cleaved bond are important for cleavage site recognition by subtilases SBT4.12, SBT4.13 and SBT5.2 (Schardon *et al*., 2016; Stintzi and Schaller, 2022). Because of its internal position in the precursor, the release of SCOOP10#1 probably requires additional step involving actions of another endoprotease and/or trimming by an exoprotease. Such complex maturation process has already been described for the maturation of the CLE19 peptide through the activity of the exoprotease Zn^2+^ carboxypeptidase SOL1 in the extracellular space (Casamitjana-Martinez *et al*., 2003; Tamaki *et al*., 2013).

Regarding the post-translational modifications, the SCOOP10 amino acid motifs containing the P hydroxylation sites were of three types: AP, TP, and GP. AP and TP are canonical motifs for the hydroxylation of P residues described for the CEP1 peptide (Ohyama *et al*., 2008) and arabinogalactan proteins (Showalter *et al*., 2010; Tan *et al*., 2003). The GP motif was found to be hydroxylated on the P residue in CLV3, CLE2 (Ohyama *et al*., 2009), and in a few other proteins (for a review see Canut *et al*., 2016). As previously reported for other cell wall proteins, the pattern of proline hydroxylation is variable (Duruflé *et al*., 2017). It has been assumed that this variability could contribute to the regulation of the biological activity or play a role in recognition of the cleavage site(s) targeted by the endoproteases as Royek *et al*. (2022) demonstrated with tyrosine sulfation. At this point, our data did not allow addressing the question of the *O*-glycosylation of the hydroxyproline residues as reported for a few CLE peptides (Shinohara and Matsubayashi, 2010, 2013; Araya *et al*., 2014; Takahashi *et al*., 2018) and PSY1 (Amano *et al*., 2007).

In our conditions, exogenous application of the synthetic SCOOP10#2 peptide, based on the native forms with or without hydroxylated prolines (GDIFTGOSGSGHGGGRTOAP or GDIFTGPSGSGHGGGRTPAP respectively), did not show any effect on seedling growth and ROS production contrary to SCOOP12 and some other members of the family. However, Rhodes *et al*. (2021) have shown that the predicted version of SCOOP10#2 peptide (FTGPSGSGHGGGR) induced a slight seedling growth inhibition and a low level of ROS at 1 μM in a MIK2 dependent manner. Hou *et al*. (2021), using another predicted sequence of the SCOOP10#2 peptide (PNGDIFTGPSGSGHGGGR, named SCOOP10#B in their publication), have shown that it induced both a strong seedling growth inhibition and ROS production at the same concentration and in a MIK2 dependent manner. These different results suggest that the SCOOP10#2 actions regarding ROS production and seedling growth inhibition might be sequence- and/or condition-dependent. The absence or the low effect of SCOOP10 peptides on ROS production could explain why the constitutive high level of transcription of *PROSCOOP10* in most of the aerial parts does not induce deleterious effects.

### SCOOP10 peptides tend to adopt a hairpin structure

The mature sequence of SCOOP10 peptides being known, we addressed the question of its molecular structure. NMR revealed that both synthetic SCOOP10#1 and SCOOP10#2 appear to be mainly unstructured in solution. Molecular dynamics simulations also pointed to the absence of a well-defined structure. Yet, the analysis of molecular dynamics trajectories showed that the peptides transiently adopt a hairpin conformation, especially in the case of SCOOP10#2. As often observed in ligand/receptor interactions, the active form of the protein might be scarcely populated and become major only in the presence of its target (Pucheta-Martinez *et al*., 2016; Sekhar *et al*., 2018). SCOOP10#2 transient structures would be stabilized by two salt-bridges. The first one, between D2 and R16 side chains, favours the formation of a turn (residues 8-9) exposing S8 and S10 while the second is between the N-terminal amine and the C-terminal carboxylate. Despite the fact that the turn is not in the middle of the structure, the two salt bridges can co-exist, because the three glycine residues in position 13-15 provide backbone flexibility. This stretch of glycines, located at the C-terminus of the S-X-S motif, is a feature shared by a majority of the SCOOP peptides and allows us to think that SCOOPs could adopt such a preferred conformation for the interaction with their receptor. Interestingly, this hairpin structure exposes the two conserved serine residues that define the SCOOP family and that were shown to be essential for the peptide function. Indeed, mutation of one of the two residues is fatal for SCOOP12 perception (Gully *et al*., 2019).

### *PROSCOOP10* delays the floral transition

The mutation of *PROSCOOP10* showed an early flowering phenotype and a lower number of leaves at bolting day compared to Col-0 plants. This observation indicates the involvement of the *PROSCOOP10* gene in flowering-time control. Multiple factors alter the flowering time such as the photoperiod, the vernalization, and the gibberellins (GAs). The pathways dependent on these factors regulate a common set of key floral integrators (Parcy *et al*., 2004; Moon *et al*., 2005; Roux *et al*., 2016; Li et al; 2010). Among them, we tested the expression of the two major genes *SOC1* and *LEAFY* (Simpson *et al*., 2002; Parcy *et al*., 2005), and of *GA3OX1-3*, involved in the GA biosynthetic pathway and promoting flowering (Blázquez *et al*., 1998). After 11 days, when the floral transition occurred (Klepikova *et al*., 2015), all three genes were up-regulated in the two early flowering *proscoop10* mutants compared to Col-0. This suggests that *PROSCOOP10* delays flowering time by repressing the expression of *SOC1, LEAFY* and *GA30X1-3*. Moreover, these results can be correlated with the *PROSCOOP10* expression profile in the aerial parts of the plant. The mining of transcriptomic data available in Genevestigator and CATdb (Zimmermann *et al*., 2004; Gagnot *et al*., 2008) revealed more information about the transcriptional regulation of *PROSCOOP10*. Indeed, in 2007, Moon *et al*. reported it as one of the only two genes significantly down regulated in the *pif1-5* mutant while describing the involvement of *PIF1* in the optimization of the greening process through the regulation of chlorophyll synthesis. Interestingly in 2018, Wu *et al*. also studied *PIF1* (called *PIL5*) and identified an early flowering phenotype under long day growth condition for the *pil5-1* mutant in which *SOC1* and *LEAFY* are also up-regulated. Moreover, Klepikova *et al*. (2015) have monitored the transcriptome of the SAM to reveal a critical time point in Arabidopsis flower initiation. In their analysis, *AT5G44580*, now identified as *PROSCOOP10* (Gully *et al*., 2019), is highly transcribed in the SAM during the first 9 days after germination, then the expression strongly decreased between 9 and 10 days, with a log2 fold change of −4.54, ranking 30 out of 968 down-regulated genes between these two stages. *PROSCOOP10* expression remains at this low level at later stages of development. This expression profile is intriguingly similar to that of *FLOWERING LOCUS* (*FLC*) another flowering regulator and opposite to *LEAFY* whose expression increase during the transition to flowering (Klepikova *et al*., 2015). While the expression of *PROSCOOP10* in leaves seems rather constitutive, its strikingly different expression profile in the SAM could fit with the early flowering phenotype of the *proscoop10* mutants. Additionally, the transcriptomic profiles of *PROSCOOP10* in various mutants or experimental conditions always showed a negative correlation between the level of *PROSCOOP10* expression and the time span before flowering (**Table 1**). Indeed, studies have shown that *PROSCOOP10* is repressed in plants showing an early flowering phenotype such as in the *abi4vtc2* and *phyABCDE* mutants compared to their corresponding wild-type (Foyer *et al*., 2012; Hu *et al*., 2013). Furthermore, in mutants such as *vtc2, tcp4* and *ga1-3* which display a late flowering phenotype or no flowering at all, *PROSCOOP10* is induced (Blazquez *et al*., 1998; Pavet *et al*., 2005; Kubota *et al*., 2017). In the *arf6-2/arf8-3* mutant, inflorescence stems elongate less than those of the wild-type and flowers arrest as infertile closed buds with short petal. In this mutant, *PROSCOOP10* is also highly expressed in comparison with the wild-type (Nagpal *et al*., 2005). All these results suggest that *PROSCOOP10* interferes with the floral transition process, upstream of *SOC1* and *LEAFY*, to delay flowering. Then, *PROSCOOP10* could be another player of the meristem identity as a delayer of floral transition, the actual signalling cascade remaining unknown. A putative role in the maintenance of the vegetative state could also be suggested after the work published by Moon *et al*. in 2007, as *PROSCOOP10* was one of the two genes co-regulated with *PIF1*, itself also reported as optimizing the greening process. The fact that *PROSCOOP10* is rather constitutively expressed in the green parts of the plant could support this hypothesis, as well as its huge decrease of expression when the SAM acquires its floral identity (Klepikova *et al*., 2015). The involvement of *PROSCOOP10* and the respective functions of both SCOOP10#1 and SCOOP10#2, that we have identified, need further investigations to clarify their roles in these mechanisms and identify their target and downstream signalling cascade. Altogether, these data may link *PROSCOO10* to the flowering-time control or the floral transition and illustrate the functional complexity of the *PROSCOOP* family.

**Table 1:** Expression profile of the *PROSCOOP10* genes extracted from public data.

| Condition | Phenotype | *PROSCOOP10* transcription | Reference |
| --- | --- | --- | --- |
| 10<sup>th</sup> vs 9<sup>th</sup> day, in SAM wild-type | wild-type | Down-regulated | Klepikova *et al.*, 2015 |
| *pif1-5* vs wild-type | early flowering | Down-regulated | Moon *et al.*, 2007 ;<br>Wu *et al.*, 2018 |
| *abi4vtc2* vs wild-type | early flowering | Down-regulated | Foyer *et al.*, 2012 |
| *phyABCDE* vs wild-type | early flowering | Down-regulated | Hu *et al.*, 2013 |
| brassinolide treatment on wild-type | early flowering | Down-regulated | Li *et al.*, 2010 |
| MeJA treatment on wild-type | early flowering | Down-regulated | Pak *et al.*, 2009 |
| 35S::*ARAF1* vs wild-type | late flowering | Up-regulated | Chavez *et al.*, 2008 |
| *vtc2* vs wild-type | late flowering | Up-regulated | Pavet *et al.*, 2005 |
| *tcp4* vs wild-type | late flowering | Up-regulated | Kubota *et al.*, 2017 |
| *ga1-3* vs wild-type | no flowering | Up-regulated | Blazquez *et al.*, 1998 |
| *arf6-2/arf8-3* vs wild-type | immature flowers | Up-regulated | Nagpal *et al.*, 2005 |

## Supplementary data

**Supplementary Table S1:** Primer sets for genotyping

**Supplementary Table S2:** ^1^H and ^13^C NMR assignment of SCOOP10#2* peptide 1 mM in 50 mM phosphate buffer (10% D_2_O), pH 6.6, 278K

**Supplementary Table S3:** ^1^H and ^13^C NMR assignment of hydroxylated SCOOP10#2 peptide

0.5 mM in 50 mM phosphate buffer (10% D_2_O), pH 6.6, 278 K

**Supplementary Table S4:** ^1^H and ^13^C NMR assignment of SCOOP10#2* peptide 0.5 mM in DMSO, 298K

**Supplementary Table S5:** ^1^H and ^13^C NMR assignment of hydroxylated SCOOP10#2 peptide

0.5 mM in DMSO, 298K

**Supplementary Table S6**: ^1^H and ^13^C NMR assignment of hydroxylated SCOOP10#1 peptide

0.5 mM in 50 mM phosphate buffer (10% D_2_O), pH 6.6, 278K

**Supplementary Table S7**: ^1^H and ^13^C NMR assignment of hydroxylated SCOOP10#1 peptide

0.5 mM in DMSO, 298K

**Supplementary Table S8:** Number of bolted inflorescences for Col-0 and *proscoop10* mutants per day after germination and per repetition

**Supplementary Figure S1: Genotyping of *proscoop10* mutant lines**

(**A**) Position of the T-DNA insertion in the *proscoop10-1* mutant. (**B**) On the left, check of the T-DNA insertion by PCR on *PROSCOOP10* and comparison with PCR on *AtCOP1*, used as a PCR positive control, in DNA of Col-0 and *proscoop10-1* mutant. On the right, check of *PROSCOOP10* expression impairment in *proscoop10-1* mutant compared to Col-0 wild-type, with *AtCOP1* used at RT-PCR positive control. (**C**) Position of the two RNA guides targeting SCOOP10#1 and SCOOP10#2 regions in the *proscoop10-2* mutant. (**D**) Reference sequence of PROSCOOP10. In red, the start and stop codons; in orange, the primer used for PCR amplification and sequencing after cloning; highlighted in grey, the guides RNAg1 and RNAg2 and in bold, the PAM1 and PAM2. (**E**) Sequencing revealed two different alleles in the therefore bi allelic *proscoop10-2* mutant. Alignments with the reference sequence shown for the first mutated allelic version a deletion of 237 bp between the two guides, in addition to nucleotide modifications in guides RNAg1 and RNAg2. This led to a deletion into SCOOP10#1 sequence and a frameshift generating a stop codon and eliminating the SCOOP10#2 sequence. For the second mutated allelic version, an important deletion of 448 bp has been generated between the two guides, leading to a break in the two sequences corresponding to SCOOP10#1 and SCOOP10#2 and a fusion of the remaining sequences. (**F**) Amino acid sequences of PROSCOOP10 in *proscoop10-2* bi allelic mutant after CRISPR/Cas9 edition. For (**A**) and (**C**) exons (CDS) and introns are represented by boxes and lines respectively. For (**A**), (**C**) and (**F**) purple: SCOOP10#1 region; blue: SCOOP10#2 region; green: signal peptide; red star: stop codon.

**Supplementary Figure S2: MS data related to SCOOP10#1 and SCOOP10#2 originating from PROSCOOP10 identified in rosette leaves apoplastic fluids**

(**A-H**) For each peptide, a representative MS/MS spectrum is shown. The positions of the hydroxyproline (O)/proline (P) residues are highlighted using the fragmentation data from the C-terminus of each peptide (y ions). The information describing the experimental procedure is available in Material and Methods.

**Supplementary Figure S3: NMR assignment and cis/trans isomerization of SCOOP10#2 in DMSO**. (**A**) NMR assignment of SCOOP10#2 reported on 1H,1H-TOCSY spectrum (HN/Hα region). (**B**) 1H,1H-NOESY cross-peaks between Hα protons of G6 and Hα or Hα protons of hydroxyproline in position 7 indicating the presence of both cis and trans conformation of the G6-P7 peptide bond, respectively.

**Supplementary Figure S4: MD simulations of SCOOP10#2 mutants**. (**A**) DSSP secondary structures calculated along the trajectories. (**B**) Contact maps of selected mutants.

**Supplementary Figure S5: Representative structures found in MD simulations for SCOOP10#2 and its mutants**. For each structure the tube (left) and ribbon (right) renderings are shown.

**Supplementary Figure S6: Structural behaviour of SCOOP10#1 in solution as monitored by NMR**. Chemical shifts deviations from random coil values of Hα protons and of the difference between Cα and Cβ carbons suggest the absence of a well definite structure for SCOOP10#1. Deviations for glycine Hα atoms were intentionally omitted.

**Supplementary Figure S7: Secondary structures and intramolecular interactions found in MD simulations of SCOOP10#1**. (**A**) DSSP secondary structures calculated along molecular dynamics (MD) simulation of hydroxylated SCOOP10#1 in solution. (**B**) Occurrence of intramolecular polar atom contacts (H-bonds and salt bridges) in SCOOP10#1 calculated along MD simulation trajectories. (**C**) Contact map.

## Acknowledgements

The authors are thankful to INRAE, CNRS, Angers University, Paul Sabatier-Toulouse 3 University, Angers Loire Métropole, French Region Pays de la Loire and Agence Nationale de la Recherche (ANR) to support their work. The authors thank Prof. Andreas Schaller (University of Hohenheim, Germany) and Prof. Cyril Zipfel (University of Zurich, Switzerland) for helpful discussions, Denis Hellal (Université Paris-Est Créteil, France) for help in the molecular dynamics simulations and Sophie Aligon (IRHS Angers, France) for plant care.

## Author contributions

SA (project coordinator), J-PR and EJ designed and supervised the experiments. M-CG, EV and MA-A performed the mutant production, phenotyping, peptide assays, and RT-qPCR analysis. TB, MZ, EJ, HC and JC performed MS analysis of apoplastic fluids. CH-L, ND’A, FR-M and ER analysed peptide structure through RMN and molecular dynamics. M-CG, ND’A, ER, J-PR, EJ and SA wrote the manuscript and all authors read and approved the final version.

## Conflict of interest

The authors declare no competing interest.

## Fundings

This research was funded by ANR (ANR-20-CE20-0025), Angers University and INRAE, and conducted in the framework of the regional program “*Objectif Végétal*, Research, Education and Innovation in Pays de la Loire”, supported by the French Region Pays de la Loire, Angers Loire Métropole and the European Regional Development Fund. Claudia Herrera-León’s PhD scholarship was funded by the National Council for Science and Technology (CONACYT).

## Data availability

All data supporting the findings of this study are available within the paper and within its supplementary materials published online. Biological materials are available from the corresponding authors upon request.

## Notes

### Competing Interest Statement

The authors have declared no competing interest.

